# Increased Excitability and Heightened Magnitude of Long-term Potentiation at Hippocampal CA3-CA1 Synapses in a Mouse Model of Neonatal Hyperoxia Exposure

**DOI:** 10.1101/2020.09.15.298513

**Authors:** Manimaran Ramani, Kiara Miller, Namasivayam Ambalavanan, Lori L McMahon

**Author notes:** **Corresponding author:** Manimaran Ramani MD, University of Alabama at Birmingham, 176F Suite 9380, 619 South 20^th^ St., Birmingham, AL 35233 USA.

## Abstract

Preterm infants exposed to supraphysiological oxygen (hyperoxia) during the neonatal period have hippocampal atrophy and cognitive dysfunction later in childhood and as adolescents. Previously, we reported that 14-week-old adult mice exposed to hyperoxia as newborns had spatial memory deficits and hippocampal shrinkage, findings that mirror those of adolescents who were born preterm. Area CA1 region of the hippocampus that is crucial for spatial learning and memory is highly vulnerable to oxidative stress. In this study, we investigated the long-term impact of neonatal hyperoxia exposure on hippocampal CA3-CA1 synaptic function. Male and female C57BL/6J mouse pups were continuously exposed to either 85% normobaric oxygen or air between postnatal days 2-14. Hippocampal slice electrophysiology at CA3-CA1 synapses was then performed at 14 weeks of age. We observed that hyperoxia exposed mice have heightened strength of basal synaptic transmission measured in input-output curves, increased fiber volley amplitude indicating increased axonal excitability, and heightened LTP magnitude at CA3-CA1 synapses, likely a consequence of increased postsynaptic depolarization during the tetanus. These data demonstrate that supraphysiological oxygen exposure during the critical neonatal developmental period leads to pathologically heightened CA3-CA1 synaptic function during early adulthood which may contribute to hippocampal shrinkage and learning and memory deficits we previously reported. Furthermore, these changes may account for cognitive disorders in children born preterm who were exposed to prolonged oxygen supplementation.

## INTRODUCTION

Children and adolescents who were born very preterm are at higher risk of a lower intelligence quotient (1) and deficits in executive function and processing speed (1, 2). They are also at high risk for developing attention deficit hyperactivity disorder and autism spectrum disorder (3, 4) than their counterparts who were born at term. In the neonatal intensive care unit, oxygen (O_2_) is the most widely used therapy. Extremely preterm infants (gestational age ≤ 28 weeks) frequently require prolonged periods of high concentrations of O_2_ due to immaturity of their lungs. Under normal conditions *in utero*, with greater adaptive mechanisms allow an adequate supply of O_2_ to the tissues, the fetus develops in a relatively hypoxemic environment (fetal oxygen saturation of 80%) (5). In contrast, for *ex utero* preterm infants, organ development occurs in a relatively hyperoxemic condition as the targeted range for O_2_ saturation (saturation range 88-95%) is higher than fetal levels, which could have a deleterious effect on organ development (6). The high concentrations of unsaturated fatty acids, high rate of O_2_ consumption, and low concentrations of endogenous antioxidants make the developing brain highly vulnerable to oxidative stress (OS) (7). Though cumulative OS has been linked with neurodegenerative disorders such as Alzheimer’s disease (8) and Parkinson’s disease (9), the impact of OS during the critical developmental period on brain development and function later in life is not well known.

Previously, we have shown that young adult C57BL/6J (14-week old) mice that are exposed to 85% O_2_ (hyperoxia) from postnatal day (P) 2 to 14 (neonatal) exhibit deficits in spatial learning along, show signs of hyperactivity and have shrinkage of area CA1 of the hippocampus (10), a brain region central to normal learning and memory (11–15) that is highly vulnerable to OS (16–19). Recently, we have shown that hyperoxia exposure during the neonatal period (P2-14) permanently impairs hippocampal mitochondrial function, increases proton leak in the mitochondria, and alters complex I enzyme function when assessed at young adult age (20). Adequate mitochondrial function is essential for normal synaptic function and long-term plasticity as well as the formation and maintenance of learning and memory. An altered hippocampal mitochondrial function could increase neuronal reactive oxygen species (ROS), disturb homeostasis, and lead to abnormalities in the induction and maintenance of hippocampal longterm potentiation (LTP), a form of long-term synaptic plasticity which results in a persistent strengthening of synaptic transmission, contributing to long-term memory formation and maintenance (21–26). While our previous research has shown that early life oxidative stress leads to deficits in cognitive-behavioral performance measured at 14 weeks of age, which are likely permanent, the mechanisms by which this occurs are still unknown.

In this study, we hypothesized that prolonged exposure to hyperoxia during the neonatal period would alter CA3-CA1 hippocampal synaptic function when assessed in early adulthood.

## MATERIALS AND METHODS

All protocols were approved by the UAB Institutional Animal Care and Use Committee (IACUC) and were consistent with the PHS Policy on Humane Care and Use of Laboratory Animals (Office of Laboratory Animal Welfare, Aug 2002) and the Guide for the Care and Use of Laboratory Animals (National Research Council, National Academy Press, 1996). Unless denoted, all experiments were done with a minimum of six mice of either sex from at least two litters for each experimental condition.

### Animal model

C57BL/6J dams and their pups were exposed to either normobaric hyperoxia (85% O_2_) or normobaric 21% O_2_ ambient air (Air) from second postnatal day (P2) until 14 days of age (P14) (27, 28). Dams were alternated every 24 hours from hyperoxia to air to reduce hyperoxia-induced adult lung injury. After the 14^th^ postnatal day (P14), mice were returned to air and maintained on standard rodent diet and light/dark cycling in microisolator cages until assessment at 14 weeks of age. A series of electrophysiological assessments were performed to study basal synaptic transmission, neurotransmitter release probability, and the induction of LTP. Animals of both sexes were used in this study.

### Hippocampal Slice Preparation

Animals were deeply anesthetized via inhalation of isoflurane, rapidly decapitated, and brains were removed. Coronal sections (350 μm) from the dorsal hippocampus were prepared using a vibratome (Leica VT10000 A). To preserve neuronal health and limit excitotoxicity, slices were sectioned in low Na^+^, sucrose-substituted ice-cold artificial cerebrospinal fluid (aCSF) (in mM: NaCl 85; KCl 2.5; MgSO_4_ 4; CaCl_2_ 0.5; NaH_2_PO_4_ 1.25; NaHCO_3_ 25; glucose 25; sucrose 75 (saturated with 95% O_2_, 5% CO_2_, pH 7.4)]. Slices were held at room temperature for 1 hr in aCSF [in mM: 119.0 NaCl, 2.5 KCl, 1.3 MgSO_4_, 2.5 CaCl_2_, 1.0 NaH_2_PO_4_, 26.0 NaHCO_3_, 11.0 Glucose [saturated with 95% O_2_, 5% CO_2_, pH 7.4]) before transfer to bath submersion chamber warmed to 26-28°C for recordings.

### Electrophysiology

As previously described (29), extracellular field excitatory postsynaptic potentials (fEPSPs) were recorded from the dendrites of CA1 pyramidal cells. CA3 axons were stimulated using a stainless-steel electrode (FHC, Bowdoin, ME, USA) placed in stratum radiatum within 200-300 μm of an aCSF-filled glass recording electrode. Baseline fEPSPs were obtained by delivering a 0.1 Hz stimulation for 200 μs to generate fEPSPs of ~0.5 mV in amplitude (~50% of maximal response). Paired-pulse facilitation (PPF) characteristic of this synapse was elicited using an inter-stimulus interval of 50 ms (30). All data were obtained using pClamp10 software (Molecular Devices, LLC, San Jose, CA.) and analyzed using GraphPad Prism 7 (GraphPad Software, Inc., and SigmaPlot V12 (Systat Software, Inc.). Only experiments with ≤8% baseline variance were included in the final data sets.

#### Input/Output Curves

Following a stable 10-minute baseline, input-output (I/O) curves were generated by increasing the stimulus intensity (10 μA increments) until a maximal fEPSP slope was obtained. The initial slopes of the six fEPSPs generated at each stimulus intensity were averaged and plotted as a single value. Fiber volley amplitudes were also measured to determine the change in axonal activation across the same stimulation range (0-320 μA). Statistical significance was determined using a two-tailed an unpaired Student’s *t*-test at the maximal stimulus intensity (320 μA; * p < 0.05).

#### Paired-Pulse Ratio (PPR)

PPR was determined by dividing the fEPSP slope of the second event in a pair of pulses at 50 ms interval by the slope of the first fEPSP. Statistical significance was determined using a two-tailed unpaired Student’s t-test at the maximal stimulus intensity (320 μA; * p < 0.05).

#### Long-term potentiation (LTP)

Following a 20 min stable baseline (0.1 Hz, 200 μs duration with the stimulus strength set to elicit initial fEPSP amplitude of ~50% maximum response), NMDA receptor (NMDAR) dependent LTP was induced using one bout of theta-burst stimulation (TBS). Each bout of TBS consists of five pulses at 100Hz repeated ten times at 200ms intervals (weak TBS). Weak TBS was chosen to ensure that differences in the LTP magnitude were not masked by using a strong TBS that could induce maximal LTP. Statistical significance was determined using a two-tailed unpaired Student’s *t*-test comparing the average of the fEPSP slope from the last 5 min of the recording (44–59 min) to baseline for the air-exposed and hyperoxia-exposed groups (*p < 0.05, **p < 0.01).

#### Postsynaptic Depolarization to Induce LTP

To determine if TBS generated a different amount of postsynaptic depolarization between experimental groups during LTP induction, the area under the curve (AUC) was measured at 100ms from baseline. This arbitrary time point was chosen due to the combined influence of NMDAR and AMPAR-mediated currents being the most impactful. Statistical significance was determined using a two-tailed unpaired Student’s *t*-test (* p < 0.05). Coastline Burst Index: As previously described (31), CA1 dendritic excitability was quantified using a modified version of the Coastline Burst Index (mCBI) (32). The equation, ∑_25_^125^(|*V*_*x*+1_ – *V_X_*|) was used to calculate mCBI, which summates the voltage differential between each point in the waveform (sampled at 10k Hz) beginning 5 ms after the stimulus artifact (25 ms). The equation also encompasses all repetitive population spikes activities (25–70 ms corresponding to 2000 total data points per waveform).

### Statistical Analysis

The experimenter was blind to the groups, data collection, and analysis. Results were expressed as means ±SE with significance set at p < 0.05 (*) determined by two-tailed unpaired Student’s *t*-test assuming unequal variance comparing results from air to hyperoxia exposed mice. Sufficient power was determined with G*Power 3.1.9.2 (Franz Faul, University Kiel, Germany). Outliers were determined with a Grubb’s test (GraphPad Software, Inc.), and significant outliers were removed.

## RESULTS

### Basal synaptic transmission is significantly altered at CA3-CA1 synapses in hyperoxia-exposed mice

To determine the lasting impact of early life O_2_ exposure on the strength of excitatory input from CA3 pyramidal cells onto CA1 pyramidal cells, we measured input-output (I/O) curves generated by incrementally increasing the stimulus intensity (0–320 μA, 10 μA intervals). As shown in **Fig 1A**, the maximum fEPSP slope is significantly larger at CA3-CA1 synapses measured in slices from the young adult mice exposed to neonatal hyperoxia compared to air-exposed mice (**Fig 1A**, at 320 μA, mean ±SE; Air = 0.31±0.06, Hyperoxia = 0.67±0.15, p = 0.04). To determine if this increase in basal synaptic strength might result from an increase in presynaptic excitability, we re-analyzed the data and plotted fiber volley amplitude, a measure of axon depolarization, versus stimulus strength. Again, we found that the maximum fiber volley amplitude at CA3-CA1 synapses was significantly larger in young adult mice exposed to neonatal hyperoxia (**Fig 1B**, at 320 μA, mean ±SE; Air = 0.75±0.19, Hyperoxia = 1.44±0.23, p = 0.04). Together, these data support the interpretation that early life O_2_ exposure increases presynaptic excitability resulting in heightened basal strength of at the CA3-CA1 synapses.

**Figure 1:**
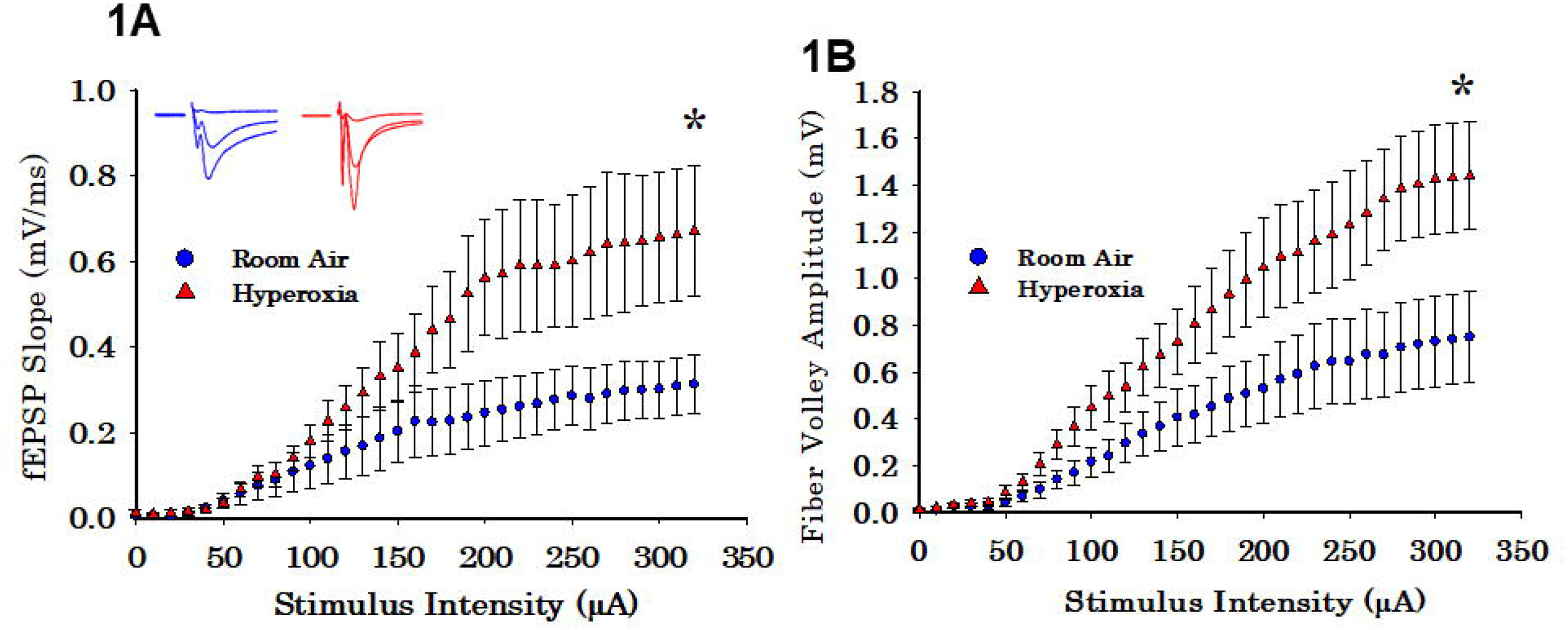
Long-term effect of neonatal hyperoxia exposure on CA3-CA1 synapses basal synaptic transmission. (A) Input-Output curves. Blue circles represent the air-exposed group, and red triangles represent the hyperoxia-exposed group. The maximum fEPSP slope is significantly larger at CA3-CA1 synapses from the young adult mice exposed to neonatal hyperoxia compared to air-exposed mice. Unpaired Student’s t-test at 320 μA means ± SEM; Air = 0.31±0.06, Hyperoxia = 0.67±0.15, p = 0.04, n= 9 slices/9 animals in Air, and 9 slices/9 animals in 85%O_2_. **\*** represents p<0.05; Air vs. Hyperoxia. (B) Fiber volley amplitude. Blue circles represent the airexposed group, and red triangles represent the hyperoxia-exposed group. Compared to air-exposed mice, fiber volley amplitude at the CA3-CA1 synapse was significantly larger in the young adult mice exposed to neonatal hyperoxia. Unpaired Student’s t-test at 320 μA; mean ±SE; Air = 0.75±0.19, Hyperoxia = 1.44±0.23, p = 0.04, n=9 slices/9 animals in Air, and 9 slices/9 animals in 85%O_2_. **\*** represents p<0.05; Air vs. Hyperoxia.

### Paired-pulse ratio is not altered in CA3-CA1 synapses in young adult mice exposed to neonatal hyperoxia

To determine whether early life O_2_ exposure alters the presynaptic release probability, we analyzed PPR, an indirect measure of presynaptic neurotransmitter release probability (33). No differences in the PPR at CA3-CA1 synapses were detectable between the adult mice that had either neonatal hyperoxia or air exposure (**Fig 2A**; mean±SE; Air = 1.48±0.08, Hyperoxia = 1.54±0.15, p = 0.75), suggesting that presynaptic release probability in adulthood is likely unaffected.

**Figure 2:**
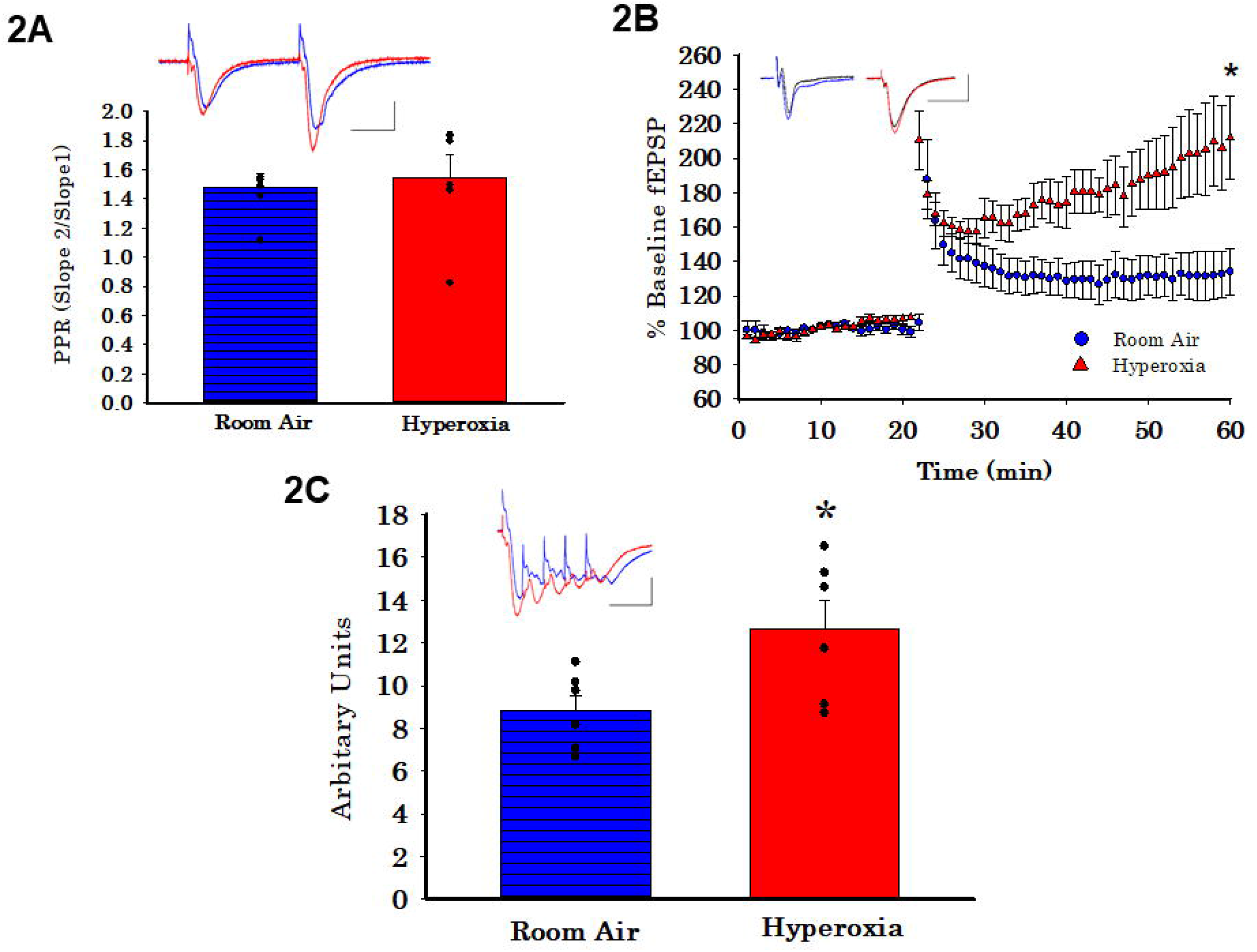
Long-term effect of neonatal hyperoxia on presynaptic probability and the magnitude of long-term potentiation in young adult mice. (A) Paired Pulse Ratio (PPR). Blue bar with horizontal lines represents air-exposed group. Solid red bar represents hyperoxia-exposed group. No differences in the PPR were detectable between the adult mice that had either neonatal hyperoxia or air exposure at the CA3-CA1 synapse. Unpaired Student’s t-test mean ±SE; Air = 1.48±0.08, Hyperoxia = 1.54±0.15, p = 0.75, n=9 slices/9 animals in Air, and 9 slices/9 animals in 85%O_2_. (B) Long-Term Potentiation (LTP). Blue circles represent the air-exposed group, and red triangles represent the hyperoxia-exposed group. The magnitude of theta-burst stimulation (TBS) induced LTP was significantly greater at the CA3-CA1 synapses in the young adult mice exposed to neonatal hyperoxia compared to air-exposed mice. Unpaired Student’s t-test: means ± SEM%; Air-exposed: 134 ± 12% of baseline fEPSP slope vs Hyperoxia-exposed 217 ± 23% of baseline fEPSP slope, n=9 slices/9 animals in Air, and 9 slices/9 animals in 85%O_2_. **\*\*** represents p < 0.01; Air vs. Hyperoxia. (C) Maximum depolarization at tetanus. Blue bar with horizontal lines represents the air-exposed group. Solid red bar represents hyperoxia-exposed group. Young adult mice that had hyperoxia exposure as neonates had a greater magnitude of the postsynaptic depolarization during the TBS. Unpaired Student’s t-test: mean ±SE; Air = 8.81±0.73, Hyperoxia = 12.66±1.34, p = 0.03, n=9 slices/9 animals in Air, and 9 slices/9 animals in 85%O_2_. **\*** represents p<0.05; Air vs. Hyperoxia.

### Heightened LTP magnitude at CA3-CA1 synapses in young adult mice exposed to neonatal hyperoxia

Next, we asked whether the increased excitability observed in the input/output curves would lead to heighten LTP magnitude since strong depolarization facilitates the removal of the voltage-dependent Mg2+ block from NMDARs. Indeed, a weak theta-burst stimulation (TBS) (1 bout of stimulation with the bout consisting of 5 pulses at 100hz) induced a greater magnitude of LTP at CA3-CA1 synapses measured at 44–59 minutes post TBS in young adult mice exposed to hyperoxia compared to air-exposed young adult mice (**Fig 2B**; Air-exposed: 134 ± 12% of baseline fEPSP slope vs. Hyperoxia-exposed 217 ± 23% of baseline fEPSP slope, **p < 0.01). Furthermore, when we measured the magnitude of the postsynaptic depolarization during the TBS, we found a larger depolarization in young adult mice exposed to hyperoxia during the neonatal period compared to air-exposed mice (**Fig 2C**; mean ±SE; Air = 8.81±0.73, hyperoxia = 12.66±1.34, p = 0.03).

### Coastline burst index indicates hyperexcitability

As a further analysis of the depolarization during TBS, we used coastline burst analysis as a measure of heightened excitability. We found that young adult mice exposed to neonatal hyperoxia had a higher mean CBI percentage compared to room-air exposed mice (**Fig 3**, mean±SE; Air = 1.06±0.03, hyperoxia = 1.27±0.05, p = 0.02). Collectively these data are consistent with the interpretation that the CA1 dendrites in young adult mice exposed to hyperoxia-during the neonatal period are hyperexcitable, perhaps a consequence of increased presynaptic excitability (**Fig 1B**) compared to air-exposed young adult mice.

**Figure 3:**
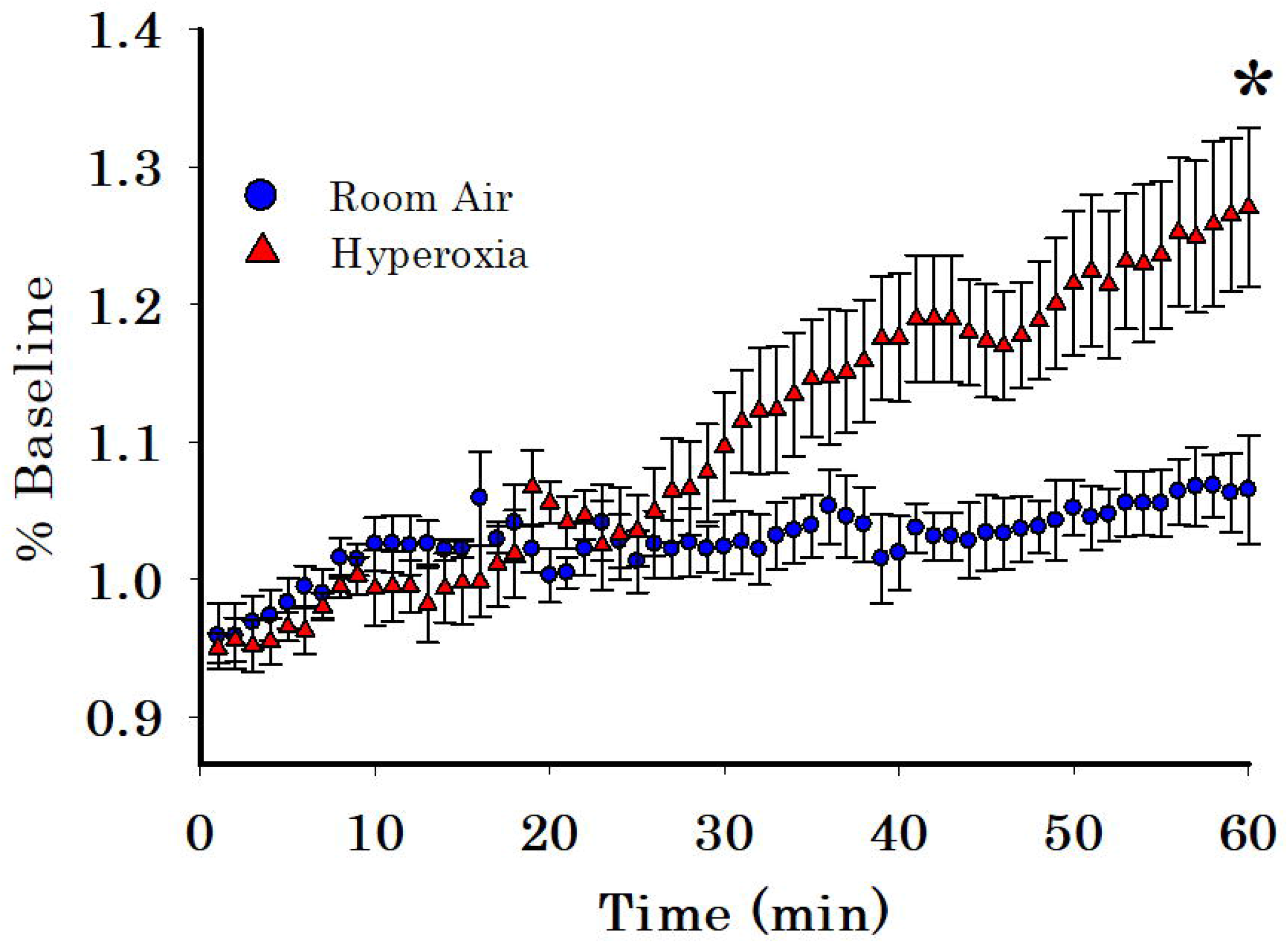
Long-term effect of neonatal hyperoxia on CA3-CA1 synapse excitability in young adult mice. Blue circles represent the air-exposed group, and red triangles represent the hyperoxia-exposed group. Young adult mice exposed to neonatal hyperoxia had higher mean costal burst index compared to air-exposed mice. Unpaired Student’s t-test: mean ±SE; Air = 1.06±0.03, Hyperoxia = 1.27±0.05, p = 0.02. **\*** represents p<0.05; Air vs. Hyperoxia.

## DISCUSSION

This is the first preclinical study to investigate the consequence of early life hyperoxia exposure, a condition experienced by preterm infants, on hippocampal synaptic function and long-term plasticity. Using acute brain slice electrophysiology in 14-week old mice, we discovered that exposure to supraphysiological levels of O_2_ during the first two postnatal weeks leads to heightened presynaptic excitability, increased strength of basal synaptic transmission, and a greater magnitude of LTP at CA3-CA1 synapses. Previously, we reported deficits in spatial memory and increased anxiety in this model (10), findings that mimic neurobehavioral deficits seen in adolescents born preterm who received prolonged periods of supplemental O_2_ in the neonatal intensive care unit. In addition to neurobehavioral deficits, preterm infants are also at high risk for clinical and subclinical seizures compared to term-born infants (22.2% compared to 0.5%) (34). Therefore, the abnormalities in synaptic transmission and heightened excitability we observe in this model suggest similar changes could be a major contributor to the cognitive and neurobehavioral deficits and seizures seen in the adolescents born preterm.

The pathophysiological processes by which early life hyperoxia exposure increases CA3-CA1 synapse excitability are likely complex. In this study, early life hyperoxia exposure leads to a larger steady-state postsynaptic dendritic depolarization later in life, which suggests that the larger magnitude of LTP we observe at CA3-CA1 synapses in the hyperoxia-exposed mice might be due to the heightened postsynaptic dendritic depolarization. Further, the increased basal synaptic strength and heightened axon excitability revealed in the I/O curves, in addition to the increase in CA1 dendritic excitability observed in the CBI analysis in the neonatal hyperoxia-exposed young adult mice, indicates that early life O_2_ exposure also markedly enhances excitability in the CA3-CA1 circuit later in life. The increased fiber volley amplitudes at CA3-CA1 synapses we observed indicate either a decrease in action potential threshold in the axons or that a larger number of axons are recruited by electrical stimulation during the generation of the I/O curves and LTP induction. The lack of change in PPR suggests there is no change in the presynaptic release probability of glutamate, indicating that the heightened responses are not likely caused by abnormal function of axon terminals. Previously, we reported that hyperoxia reduces the expression of hippocampal potassium voltage-gated channel subfamily D member 2 (KCND2) protein at 14 weeks (35). Therefore it is possible that the action potential threshold is decreased as a consequence of neonatal hyperoxia. Clearly, future studies are needed to determine how hyperoxia conditions during the neonatal period induce altered excitability and whether axons sprouting occurs that remains in adulthood.

Another important consideration is the possibility of decreased GABAergic inhibition, as inhibitory interneurons play a critical role in maintaining the excitatory and inhibitory balance. Since developing GABAergic interneurons are highly vulnerable to oxidative stress (36), it is highly likely that hyperoxia-exposed young-adult mice may also have deficits in GABAergic function. Thus, weakened inhibition would be expected to facilitate the induction of LTP, leading to a heightened potentiation, and it could also be responsible for the CA1 dendritic hyperexcitability we observed in our neonatal hyperoxia model. In future studies, detailed investigations are needed into a potential loss of GABAergic interneurons leading to decreased strength of inhibitory transmission and increased the excitation/inhibition balance.

Our study used TBS to induce LTP (37) instead of high-frequency stimulation. TBS mimics the complex-spike discharges of the pyramidal neurons (38, 39) that occur spontaneously during learning and memory formation. The optimal repetition rate used in TBS corresponds to the frequency of the hippocampal theta rhythm, an EEG pattern related to memory storage processes (40). Though this study assessed the changes in LTP magnitude at CA3-CA1 synapses induced by early life OS, we did not investigate possible changes in LTP induction or expression at other hippocampal synapses such as the perforant pathway to the dentate gyrus or to distal CA1 dendrites (temporammonic pathway), which provide major input to the hippocampus from the entorhinal cortex and is involved in memory consolidation.

Our findings are consistent with neonatal mouse models of ischemia and reperfusion injury (41) but are in contrast with many models of OS-associated brain disorders, which linked OS-induced LTP deficits to the cognitive decline (42–44). This indicates that OS affects the developing brain differently than the adult brain and leads to long-lasting impairment in synaptic function. Additional electrophysiological studies are needed to assess the excitatory and inhibitory balance, action potential generation, interneuron function, axonal sprouting, and dendritic arborization. A better understanding of these systems is required to determine the mechanisms by which early life OS leads to long-lasting synaptic dysfunction.

## Acknowledgments

This work was partially funded by R01AG021612, R01NS076312, R25NS089463, a KPRI grant, and the Division of Neonatology, Department of Pediatrics, University of Alabama at Birmingham, Alabama USA.

## Author contributions

Concept and design: MR and LLM; Electrophysiology: KLM. Data analysis and interpretation: MR, NA, KLM and LLM; Drafting the manuscript for important intellectual content: MR, KLM, NA, and LLM. All authors edited and approved the final version of the manuscript.

